# Linkage disequilibrium and haplotype block patterns in popcorn populations

**DOI:** 10.1101/688960

**Authors:** Andréa Carla Bastos Andrade, José Marcelo Soriano Viana, Helcio Duarte Pereira, Vitor Batista Pinto, Fabyano Fonseca e Silva

## Abstract

Linkage disequilibrium (LD) analysis provides information on evolutionary aspects of the populations and allows selecting populations and single nucleotide polymorphisms (SNPs) for association studies. Recently, haplotype blocks have been used to increase the power of quantitative trait loci detection in genome-wide association studies and the prediction accuracy with genomic selection. The objectives of this study were to compare the degree of LD, the LD decay, the LD decay extent, and the number and length of haplotype blocks in the populations and to elaborate the first LD map for maize, for elucidating if the maize chromosomes also had a pattern of interspaced regions of high and low rates of recombination. We used a biparental temperate population, a tropical synthetic, and a tropical breeding population, genotyped for approximately 75,000 SNPs. The level of LD expressed by the r^2^ values is surprisingly low (0.02, 0.04, and 0.04), but comparable to some non-isolated human populations. The general evidence is that the synthetic is the population with higher LD. It is not expected a significant advantage of haplotype-based association study and along generations genomic selection due to the reduced number of SNPs in the haplotype blocks (2 to 3). The results concerning LD decay (rapid decay after 5-10 kb) and LD decay extent (along up to 300 kb) are in the range observed with maize inbred line panels. Our most important result is that maize chromosomes had a pattern of regions of extensive LD interspaced with regions of low LD. However, our simple simulated LD map provides evidence that this pattern can reflect regions with differences of allele frequencies and LD level (expressed by D’) and not regions with high and low rates of recombination.

## Introduction

Linkage disequilibrium (LD) analysis is important to humans, other animal species, and plants because the results can be used for positional cloning, provide information on rate of recombination, gene conversion, and evolutionary aspects of the populations, including recombination history, mutation, selection, genetic drift, and admixture, and allows selecting populations and single nucleotide polymorphisms (SNPs) for association studies [1]. The most common LD measures are D’ and r^2^. The statistic D’ is the ratio between D (the difference between products of haplotypes, D = P(AB).P(ab) – P(Ab).P(aB)) and the deviation of the actual gametic frequency from linkage equilibrium [2]. The statistic r^2^ is the square of the correlation between the values of alleles at two loci in the same gamete, where D is the covariance [3].

Additional information on historical recombination is provided by the analysis of the haplotype blocks pattern in populations. A haplotype block is a chromosome region in which there are few haplotypes (combinations of alleles of multiple SNPs within a haplotype block) (2–4 per block), and for which the LD analysis provides evidence of a low rate of recombination [1]. Recently, haplotype blocks have been used to increase the power of QTL (quantitative trait loci) detection in genome-wide association studies (GWAS) and the prediction accuracy with genomic selection. Based on a panel including 183 maize inbred lines genotyped for 38,000 SNPs, Maldonado, Mora (4) confirmed the advantage of haplotype-based GWAS for ear and plant height, ratio ear height/plant height, and leaf angle, compared with the single SNP analysis. Hess, Druet (5) observed an increase of up to 5.5% in the accuracy of genomic prediction in an admixed dairy cattle population using fixed-length haplotypes, relative to the single SNP approach. Although there are several methods for defining haplotype blocks, the most common procedure was proposed by Gabriel, Schaffner (6). Their criterion is that the one-sided upper 95% confidence bound on D’ is > 0.98 and the lower bound is > 0.70.

The characterization of the LD and haplotype block patterns in human, domesticated animal, and plant populations have provided variable results concerning the degree of LD, LD decay, LD decay extent, and number and length of the haplotype blocks. Most maize LD studies have been done with inbred line panels. Thirunavukkarasu, Hossain (7) and Truntzler, Ranc (8) observed an overall average r^2^ between 0.23 and 0.61, LD decay after 5-10 kb, and LD extent along 200-300 kb. Faster LD decay and shorter LD extent (less than 4 kb) were observed by Maldonado, Mora (4). Higher LD and slower LD decay was observed in biparental and multiparental maize populations [9]. The number and length of haplotype blocks is also highly variable [4, 7].

In several investigations in human populations the structure of LD was described based on LD maps. In an LD map, each SNP has a LD position in LD units (LDUs). One LDU is the distance in kilobases at which disequilibrium (expressed as the Malecot’s prediction of association - ρ) declines to approximately 0.37 of its starting value. Assuming unrelated individuals, ρ equates to the absolute value of D’. The difference between the LD positions of two SNPs divided by the distance in kilobases (d) is the exponential decline of disequilibrium (ε). LDUs share an inverse relationship with the recombination rate. Thus, regions with extensive disequilibrium have few LDUs (plateaus or blocks) and regions with many LDUs have high levels of recombination rate (steps). Holes in the LD maps are regions where greater marker density is required to provide a full characterization of the block and step patterns of the LD. Holes are identified by a LD map interval of 3, which is an arbitrary value because disequilibrium is indeterminate for εd > 3 and of doubtful reliability for εd > 2 [10, 11].

Because there is no information on LD and structure of haplotype blocks in popcorn populations and no LD maps for maize, the objectives of this study were to compare the degree of LD, the LD decay, the LD decay extent, and the number and length of haplotype blocks in the populations and to elaborate the first LD map for maize, for elucidating if the maize chromosomes also had a pattern of interspaced regions of high and low rates of recombination.

## Materials and Methods

### Populations

We used a biparental (F_2_ generation) temperate population, a tropical synthetic (Synthetic UFV), and a tropical breeding population (Beija-Flor cycle 4). The biparental population was derived from the single cross AP4502, developed by Agricultural Alumni Seed Improvement Association, Romney, IN, USA. The Synthetic UFV and Beija-Flor cycle 4 (BFc4) were developed by Federal University of Viçosa (UFV), Minas Gerais, Brazil. The synthetic was derived by random crossings involving 20 elite inbred lines from the tropical population Viçosa and 20 elite inbred lines from the tropical population Beija-Flor. The inbred lines were selected based on expansion volume (a measure of popcorn quality). Beija-Flor cycle 4 was developed after four cycles of half-sib selection based on expansion volume. Theoretically, a biparental population shows LD only for linked genes and molecular markers. A synthetic there is LD for genes and molecular markers with independent assortment.

### DNA extraction, genotyping-by-sequencing (GBS), SNP calling, data quality control, and imputation

Leaf samples of young plants were collected for DNA extraction. The DNA extraction was performed using the CTAB (cetyl trimethylammonium bromide) protocol with modifications. After quantification, the DNA samples of 574 plants (190 or 192 from each population) were sent to the Institute of Biotechnology at Cornell University (two plates of 95 samples from the biparental population) and Institut de Recherche en Immunologie et en Cancérologie/IRIC at University of Montreal (four plates of 96 samples from the tropical populations) for GBS services based on HiSeq 2500 and NextSeq500, respectively. The SNP variant call services were provided by the Institute of Biotechnology and Omega Bioservices, Norcross, GA, respectively, using B73 version 4 as the reference genome. After reading the data using the R package vcfR [12], we filtered by missing allele and chromosome. Then, we computed the SNP and genotype call rates and the minor allele frequency (MAF), employing the R package HapEstXXR [13]. After filtering by MAF > 0.01, we imputed based on Beagle [14], using the R package synbreed [15]. The number of SNPs after the data quality control and imputation were 145,420, 74,773, and 76,055 for the biparental population, Synthetic UFV, and Beija-Flor c4, respectively. To maintain a similar number of SNPs for the populations, we finally performed a random sampling of 75,000 SNPs from the biparental population.

### LD and haplotype block analyses

For the Hardy-Weinberg equilibrium analysis by population and chromosome it was adopted the Bonferroni criterion to keep a global level of significance of 1%. To characterize the block and step patterns of LD in the populations we constructed the LD maps by chromosome using the interval method [16]. To evaluate if the LD maps allow inference on the overall degree of LD by chromosome in the populations we also processed a simulated data set, generated with *REALbreeding* software (available by request). This software has been recently used in studies of population structure [17], QTL mapping [18], genomic selection [19], and genome-wide association studies [20]. We simulated the genotyping of 200 individuals in a population (generation 0) and 200 individuals in the same population after 10 generations of random crossings (generation 10), for 287 SNPs covering 298 cM (density of 1 cM) of a single chromosome.

We then evaluated the degree of LD by chromosome in the populations concerning SNPs separated by up to 500 kb, using a two marker expectation-maximization (EM) algorithm [21]. The LD analyses were based on the D’ absolute value (|D’|) and r^2^. The physical distances between SNPs were classified into six intervals of 50 kb (0-50 to 451-500) to study the LD decay and LD decay extent. To define a haplotype block, we adopted the criterion proposed by Gabriel, Schaffner (6). The haplotypes were estimated using an accelerated EM algorithm with a partition-ligation approach [22], to generate phased haplotypes for population frequency [23].

The LD and haplotype block analyses were also performed at the intragenic level. We choose 12 genes related to zein (one), starch (four), cellulose (five), and fatty acids biosynthesis (two) (S1 Table). With two exceptions, the selected genes had at least five SNPs in each population (maximum of 21). For the intragenic LD decay and LD decay extent analyses we computed the average |D’| and r^2^ values defining intervals of 1 kb (0-1 to 10.1-11 kb). All analyses were performed using LDMAP [16] and Haploview [21]. To assess the haplotype blocks information, the haplotype files for each population and chromosome were read by a program (*Haplotype blocks summary*) developed in REALbasic 2009 by Prof. José Marcelo Soriano Viana.

## Results

Versus the physical maize map (available at https://www.maizegdb.org/), the GBS provided a SNP coverage between 99.5 to practically 100.0% of the genomes of all chromosomes, in each population (Table 1). Except chromosome 10 in the breeding population, the number of SNPs was generally in proportion to the chromosome length, providing a SNP density in the range 23.5 to 44.3 kb (one SNP per 30.0 kb on average). The average MAF was approximately 0.1 regardless of chromosome and population but the populations differ in the MAF distribution. The biparental population has a bimodal distribution and shows the higher number of SNPs with frequencies close to 0.01 and greater than 0.45 (S2 Figure). The synthetic and the breeding population have similar MAF distributions. The analysis of Hardy-Weinberg equilibrium evidenced that most of the SNPs in the biparental population had a non significant deviation whereas most of the SNPs in the other populations showed a significant deviation. We retained SNPs with significant deviation from the Hardy-Weinberg equilibrium in the synthetic and breeding population to keep a similar number of SNPs for the LD and haplotype block analyses. To maintain a similar number of SNPs for constructing the LD maps by chromosome, we used the SNPs in Hardy-Weinberg equilibrium in the synthetic and breeding population as well as a sample of SNPs with no significant deviation from the Hardy-Weinberg equilibrium from the biparental population.

**Table 1.** Number of SNPs, SNP coverage (kb), average SNP interval (bp), MAF, and minimum, average, and maximum LD measures by chromosome in each population.

| Population | Chr. | SNPs | SNP coverage | SNP interval | MAF | \|D'\| |  |  | r<sup>2</sup> |  |  |
| --- | --- | --- | --- | --- | --- | --- | --- | --- | --- | --- | --- |
|  |  |  |  |  |  | Min. | Av. | Max. | Min. | Av. | Max. |
| Biparental | 1 | 11,816 | 307,039.27 | 25,982.75 | 0.09 | 0.0 | 0.78 | 1.0 | 0.00 | 0.023 | 1.00 |
|  | 2 | 8,710 | 244,412.25 | 28,059.68 | 0.11 | 0.0 | 0.77 | 1.0 | 0.00 | 0.026 | 1.00 |
|  | 3 | 8,205 | 235,520.19 | 28,693.18 | 0.11 | 0.0 | 0.75 | 1.0 | 0.00 | 0.032 | 1.00 |
|  | 4 | 8,081 | 246,827.22 | 30,525.85 | 0.07 | 0.0 | 0.81 | 1.0 | 0.00 | 0.015 | 1.00 |
|  | 5 | 8,697 | 223,657.67 | 25,708.94 | 0.09 | 0.0 | 0.79 | 1.0 | 0.00 | 0.019 | 1.00 |
|  | 6 | 5,883 | 173,906.61 | 29,537.18 | 0.10 | 0.0 | 0.78 | 1.0 | 0.00 | 0.027 | 1.00 |
|  | 7 | 6,401 | 182,200.48 | 28,440.64 | 0.11 | 0.0 | 0.77 | 1.0 | 0.00 | 0.025 | 1.00 |
|  | 8 | 6,528 | 181,042.64 | 27,725.54 | 0.10 | 0.0 | 0.78 | 1.0 | 0.00 | 0.023 | 1.00 |
|  | 9 | 5,625 | 159,429.26 | 28,336.11 | 0.11 | 0.0 | 0.76 | 1.0 | 0.00 | 0.027 | 1.00 |
|  | 10 | 5,054 | 150,832.73 | 29,824.39 | 0.10 | 0.0 | 0.76 | 1.0 | 0.00 | 0.025 | 1.00 |
| Synthetic | 1 | 11,224 | 306,909.66 | 27,341.76 | 0.10 | 0.0 | 0.75 | 1.0 | 0.02 | 0.046 | 1.00 |
|  | 2 | 9,712 | 244,369.34 | 25,159.97 | 0.10 | 0.0 | 0.75 | 1.0 | 0.02 | 0.041 | 1.00 |
|  | 3 | 9,374 | 235,478.72 | 25,083.00 | 0.10 | 0.0 | 0.76 | 1.0 | 0.02 | 0.042 | 1.00 |
|  | 4 | 5,840 | 246,943.47 | 42,170.02 | 0.10 | 0.0 | 0.74 | 1.0 | 0.02 | 0.052 | 1.00 |
|  | 5 | 9,460 | 223,706.51 | 23,589.54 | 0.10 | 0.0 | 0.74 | 1.0 | 0.02 | 0.040 | 1.00 |
|  | 6 | 5,294 | 173,221.42 | 32,692.62 | 0.10 | 0.0 | 0.74 | 1.0 | 0.02 | 0.050 | 1.00 |
|  | 7 | 6,299 | 182,159.80 | 28,857.92 | 0.11 | 0.0 | 0.74 | 1.0 | 0.02 | 0.042 | 1.00 |
|  | 8 | 6,248 | 180,660.38 | 28,850.52 | 0.10 | 0.0 | 0.76 | 1.0 | 0.02 | 0.044 | 1.00 |
|  | 9 | 5,161 | 159,553.33 | 30,909.31 | 0.11 | 0.0 | 0.75 | 1.0 | 0.02 | 0.045 | 1.00 |
|  | 10 | 6,161 | 150,828.61 | 24,464.31 | 0.09 | 0.0 | 0.75 | 1.0 | 0.02 | 0.034 | 1.00 |
| BFc4 | 1 | 10,182 | 306,774.01 | 30,126.80 | 0.11 | 0.20 | 0.71 | 1.0 | 0.02 | 0.047 | 1.00 |
|  | 2 | 8,481 | 244,407.97 | 28,816.88 | 0.11 | 0.21 | 0.69 | 1.0 | 0.02 | 0.042 | 1.00 |
|  | 3 | 8,005 | 235,478.74 | 29,373.18 | 0.11 | 0.20 | 0.70 | 1.0 | 0.02 | 0.040 | 1.00 |
|  | 4 | 5,558 | 246,840.44 | 44,379.59 | 0.11 | 0.20 | 0.69 | 1.0 | 0.02 | 0.054 | 1.00 |
|  | 5 | 7,674 | 223,706.51 | 29,080.32 | 0.11 | 0.20 | 0.70 | 1.0 | 0.02 | 0.039 | 1.00 |
|  | 6 | 4,547 | 173,351.50 | 38,093.29 | 0.11 | 0.19 | 0.68 | 1.0 | 0.02 | 0.044 | 1.00 |
|  | 7 | 5,602 | 182,155.19 | 32,448.24 | 0.11 | 0.20 | 0.69 | 1.0 | 0.02 | 0.040 | 1.00 |
|  | 8 | 5,020 | 180,660.38 | 35,943.93 | 0.12 | 0.20 | 0.70 | 1.0 | 0.02 | 0.048 | 1.00 |
|  | 9 | 5,353 | 159,489.87 | 29,788.60 | 0.11 | 0.20 | 0.69 | 1.0 | 0.02 | 0.042 | 1.00 |
|  | 10 | 15,633 | 150,926.35 | 9,653.39 | 0.13 | 0.20 | 0.52 | 1.0 | 0.02 | 0.021 | 1.00 |

The LD map from the simulated data evidence that the LD units were lower for the generation with lower LD (generation 10) (Figure 1). Thus, the LD maps by chromosome reveal that the higher global LD (in LDU) was observed in the synthetic but only for chromosomes 1 to 7 (S3 Figure). The higher global LD for chromosomes 8 and 9 was observed in the biparental population. The higher global LD for chromosome 10 was seen in the breeding population. The lower global LD was observed in chromosome 6 and the higher global LD was observed in chromosome 10 of the breeding population. Because of the much higher number of SNPs in Hardy-Weinberg equilibrium in the biparental population, we only used this population for analysis of the number and length of the hot (high recombination rate) and cold (low recombination rate) spots regions of the chromosomes, as well as the number and length of the holes (Table 2). Except for chromosome 10, where the average lengths of the hot and cold spots regions were approximately 37 and 38 kb, respectively, the average lengths of the hot and cold spots regions for the other chromosomes ranged between approximately 45-55 and 83-110 kb, respectively. The number of hot spots ranged between 1,788 and 3,897 and the number of cold spots ranged from 608 to 1,507. The holes represented only 0.4 to 2.7% of the chromosomal genomes.

**Figure 1.**
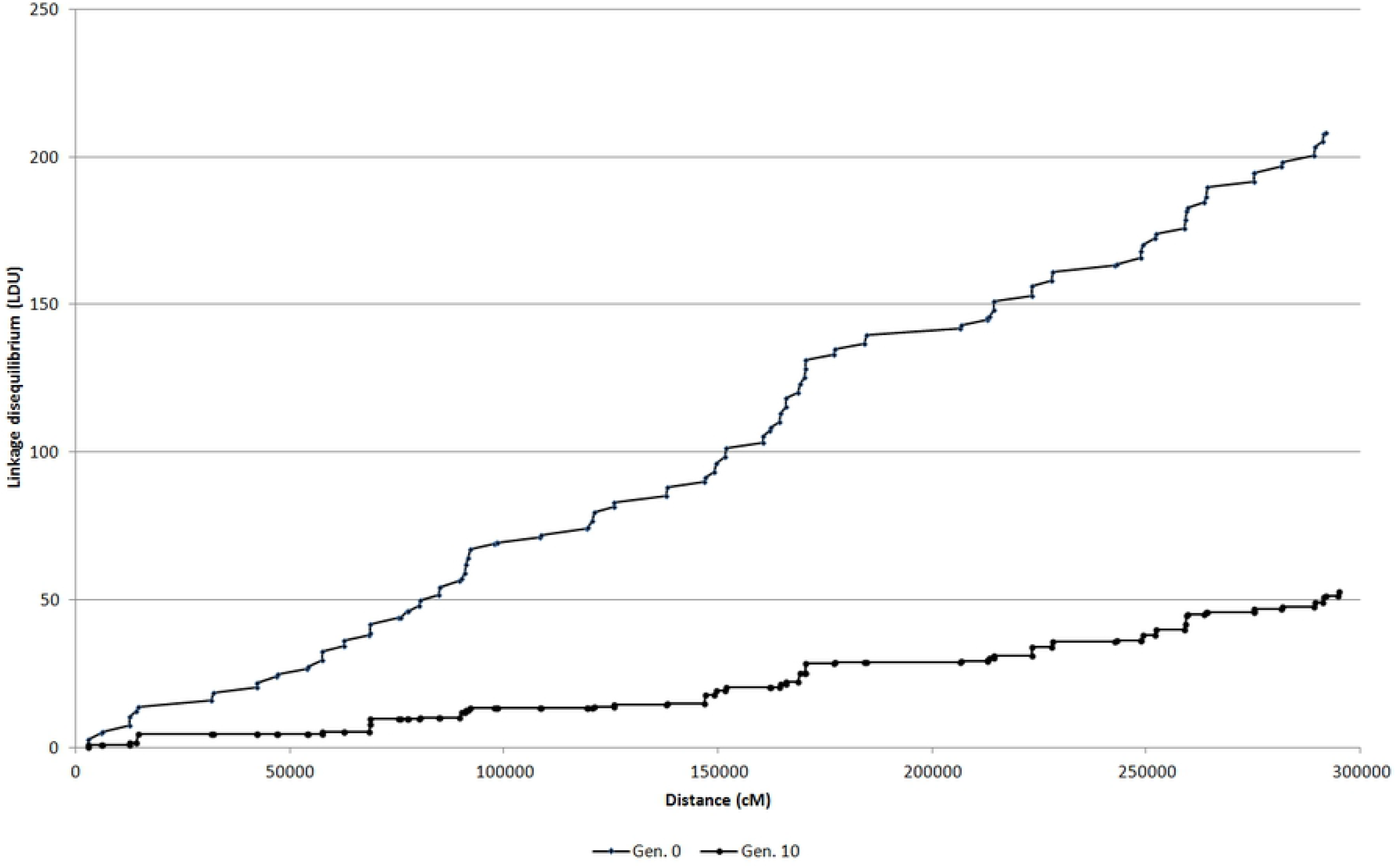
LD maps for generations 0 and 10.

**Table 2.**
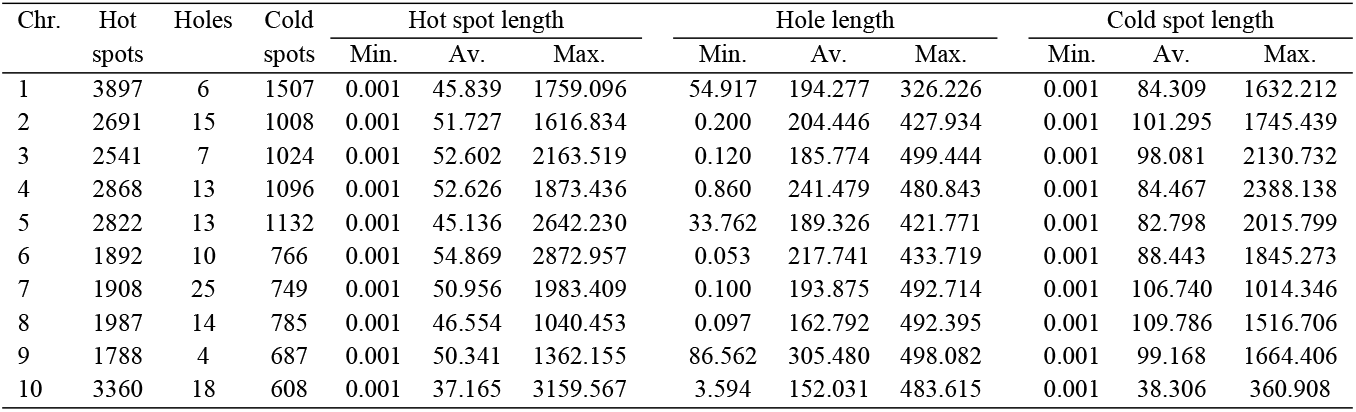
Number and minimum, average, and maximum length (kb) of the hot spots (steps), holes, and cold spots (plateaus) by chromosome in the biparental population.

Concerning SNPs separated by up to 500 kb, the biparental population and the synthetic have similar average |D’| values (0.77 and 0.75). The values are approximately 10-14% greater than the average value in the breeding population (Table 1). Interestingly, the average r^2^ value in the biparental population is approximately half of the corresponding average values observed in the other populations (0.02 versus 0.04, and 0.04). Regardless of the chromosome, the populations with the higher and lower frequencies of |D’| values greater than 0.75 are the biparental (65-74%) and the breeding population (26-58%), respectively. However, the frequency of r^2^ values greater than 0.75 is lower in the biparental population (0.2-0.5%) and higher in the other populations (0.2-1.6%) (S4 Table). Furthermore, the average distance for SNPs with r^2^ values greater than 0.75 are much higher in the biparental population (approximately 80 to 126 kb). In the other populations the ranges are approximately 6 to 19 and 6 to 35 kb. There are no differences between the populations regarding the average distance for SNPs with |D’| values greater than 0.75 (in the range of approximately 207 to 229 kb).

Regardless of the chromosome, population, and LD measurement, the LD decreased as the between-SNP distance increased from 0-50 to 451-500 kb (S5 and S6 Figures). In general, there is an initially higher LD decrease for SNPs separated by 51-100 kb (3 to 7% for |D’| and 28 to 66% for r^2^, on average) and then a gradual decrease to the minimum LD value for SNPs separated by 451-500 kb. Because there are no significant differences between chromosomes, we can state that following an initial higher decrease after 50 kb the |D’| and the r^2^ in the biparental population extends with similar magnitude for an interval of 450 kb (Figure 2). In this interval, the average |D’| values decreased from 0.69-0.77 to 0.64-0.77 in the three populations and the average r^2^ values in the biparental population decreased from 0.025 to 0.020. However, in the other two populations the average r^2^ value decreased in approximately 50%. The r^2^ decay from its maximum average value reached 36 to 73% after 5-10 kb (Figure 2c).

**Figure 2.**
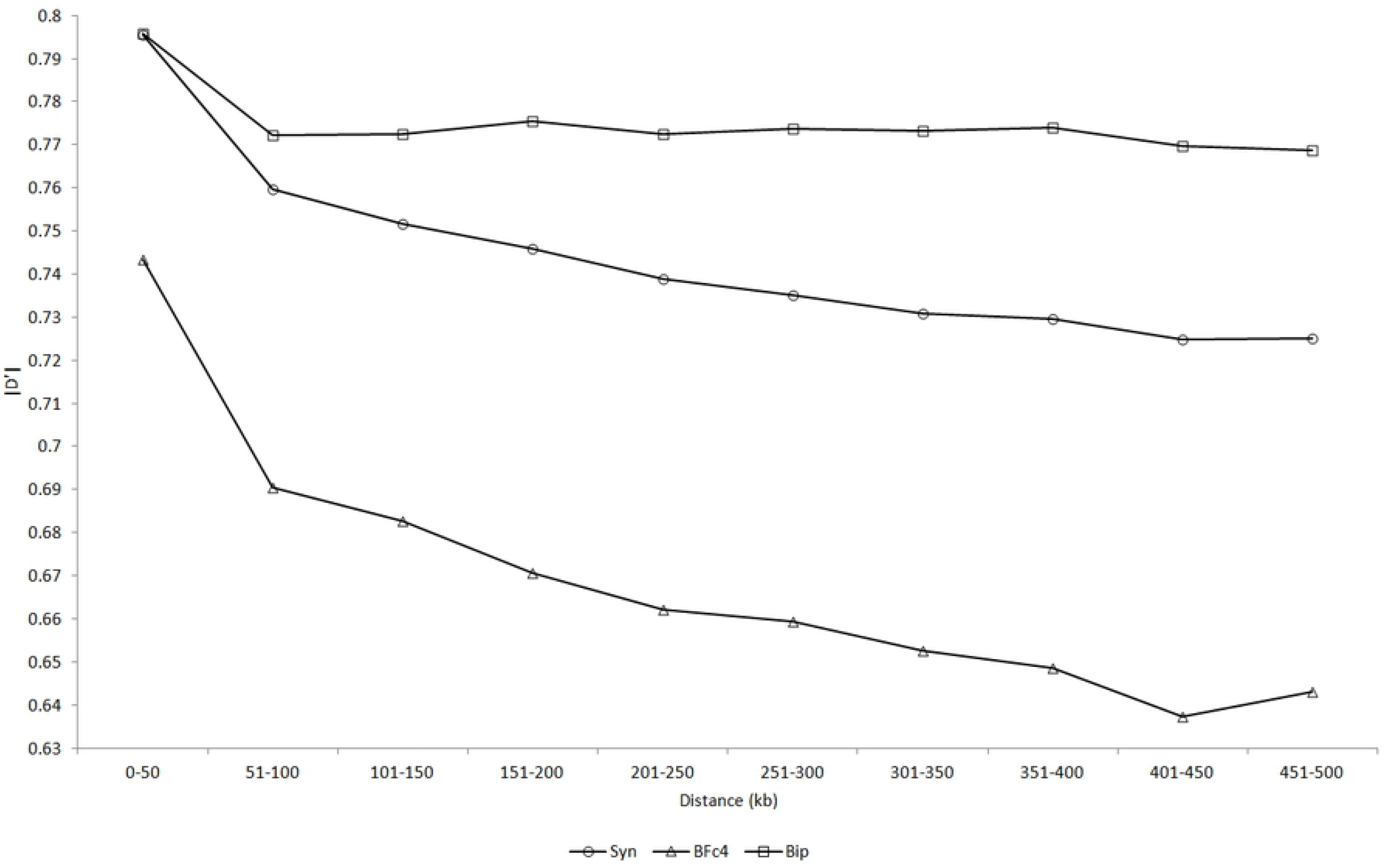
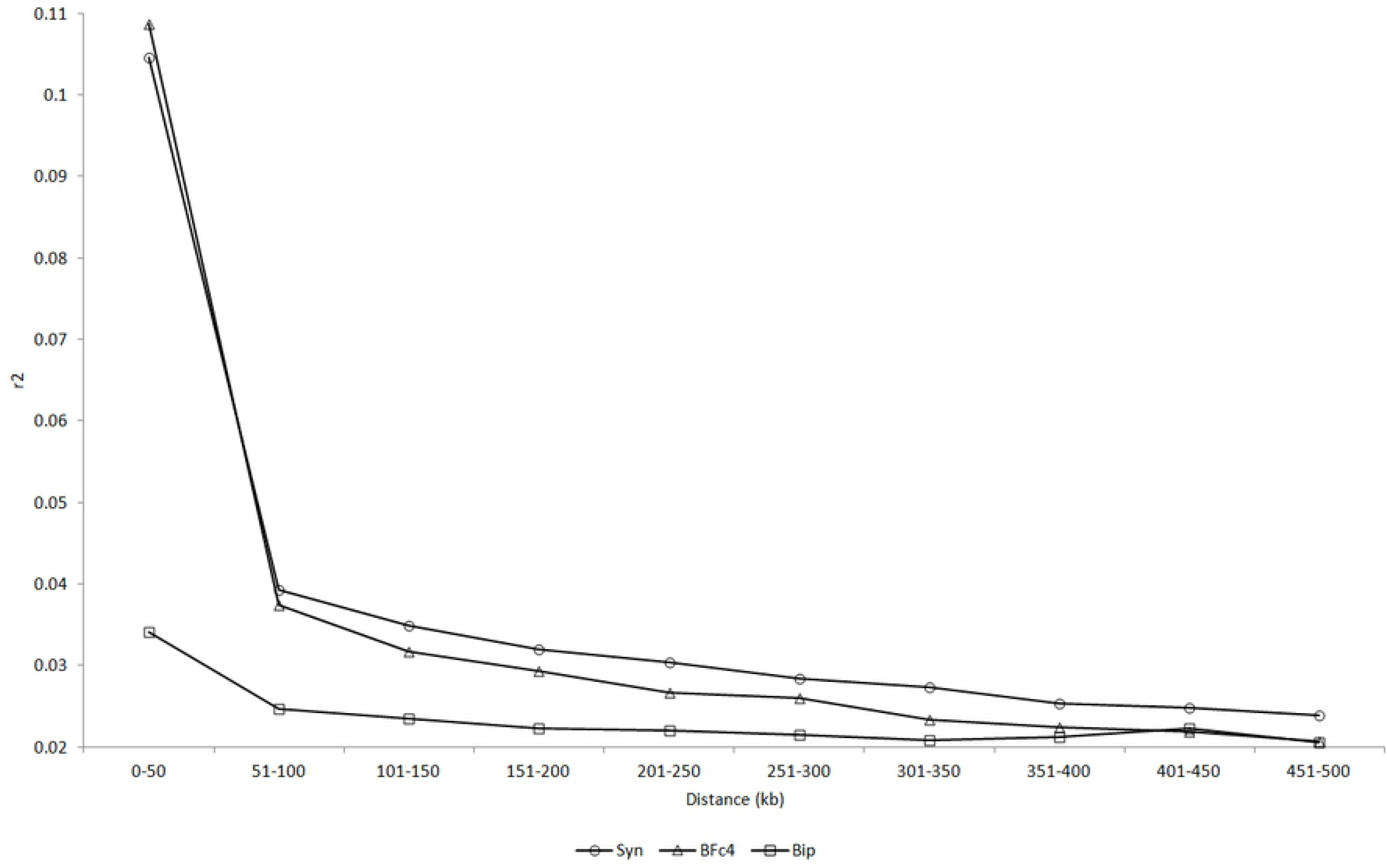
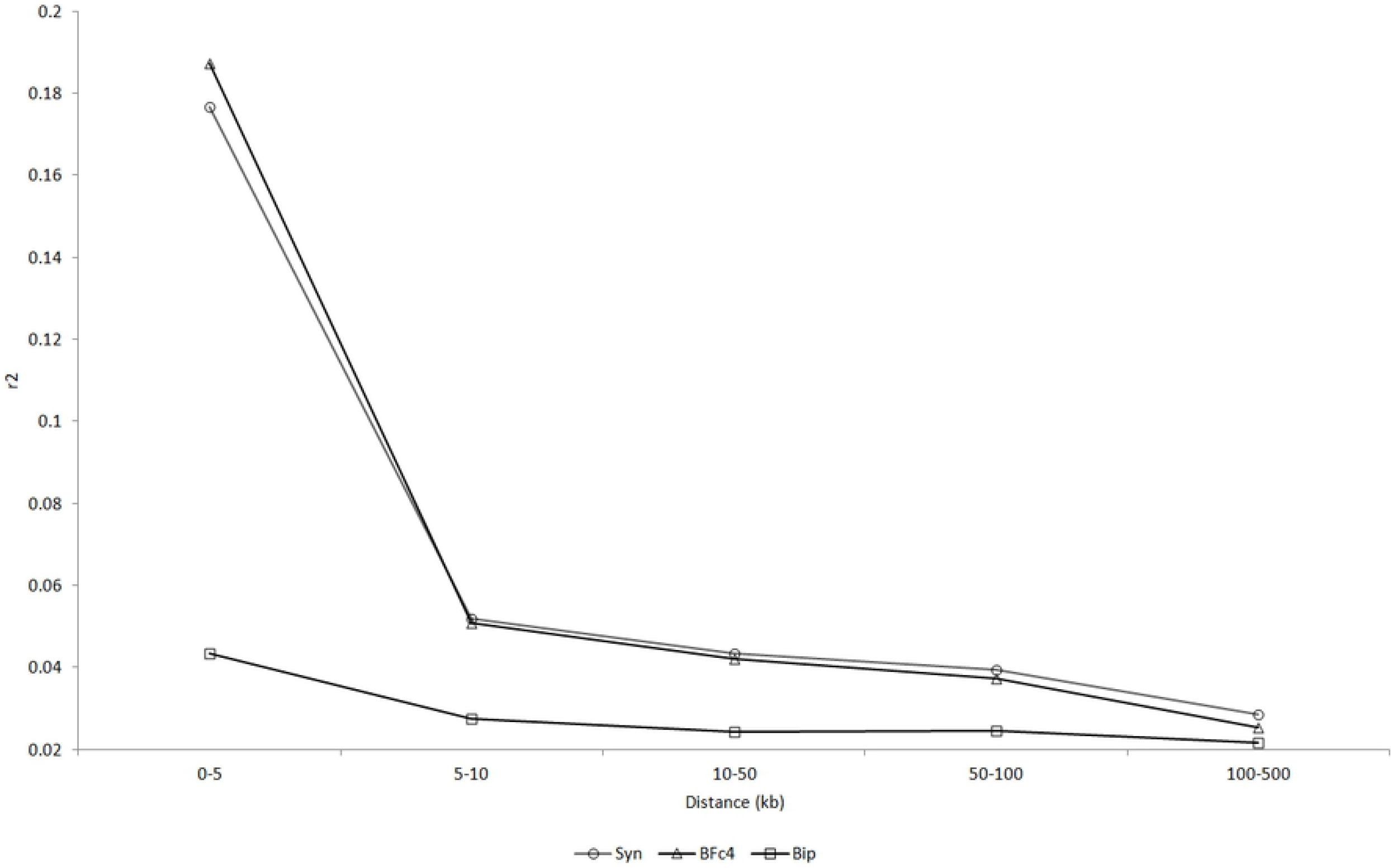
Overall average |D’| (a) and r^2^ (b and c) values by distance interval (kb) in the biparental population (Bip), in the synthetic (Syn), and in the breeding population (BFc4).

The biparental population also differs from the other populations concerning the pattern of haplotype blocks (Table 3). The biparental population presents the lower average number of haplotype blocks per chromosome (approximately 225 versus 700 and 730 on average), the lower block length (approximately 1 versus 11 kb on average), and the lower number of SNPs per block (approximately 2 versus 3 on average). Most of the haplotype blocks in the three populations include two SNPs but the number of haplotype blocks with three or more SNPs is greater in the synthetic and breeding population (S7 Figure). It is important to highlight that the total length of the haplotype blocks represents only 0.01 to 5.13% of the chromosome genomes.

**Table 3.** Haplotype blocks structure of the populations.

| Population | Chr. | Blocks | Block size (kb) |  |  |  | SNPs |  |  |  |
| --- | --- | --- | --- | --- | --- | --- | --- | --- | --- | --- |
|  |  |  | Total | Mean | Min. | Max. | Total | Mean | Min. | Max. |
| Biparental | 1 | 336 | 58.60 | 0.17 | 0.001 | 10.30 | 727 | 2.2 | 2 | 5 |
|  | 2 | 294 | 588.31 | 2.00 | 0.001 | 298.9 | 647 | 2.2 | 2 | 6 |
|  | 3 | 273 | 307.66 | 1.13 | 0.001 | 101.90 | 622 | 2.3 | 2 | 5 |
|  | 4 | 193 | 35.80 | 0.19 | 0.001 | 23.15 | 430 | 2.2 | 2 | 6 |
|  | 5 | 218 | 47.49 | 0.22 | 0.001 | 20.39 | 484 | 2.2 | 2 | 4 |
|  | 6 | 169 | 419.24 | 2.48 | 0.001 | 292.35 | 387 | 2.3 | 2 | 5 |
|  | 7 | 215 | 45.60 | 0.21 | 0.001 | 11.68 | 479 | 2.2 | 2 | 5 |
|  | 8 | 186 | 511.79 | 2.75 | 0.001 | 423.79 | 409 | 2.2 | 2 | 5 |
|  | 9 | 195 | 58.19 | 0.29 | 0.001 | 15.58 | 432 | 2.2 | 2 | 5 |
|  | 10 | 170 | 314.88 | 1.85 | 0.001 | 307.49 | 370 | 2.2 | 2 | 4 |
| Synthetic | 1 | 1126 | 11935.23 | 10.60 | 0.001 | 494.94 | 3093 | 2.7 | 2 | 10 |
|  | 2 | 935 | 8501.15 | 9.09 | 0.001 | 451.74 | 2565 | 2.7 | 2 | 11 |
|  | 3 | 810 | 9065.75 | 11.19 | 0.001 | 457.30 | 2257 | 2.8 | 2 | 11 |
|  | 4 | 525 | 6615.63 | 12.60 | 0.001 | 423.71 | 1409 | 2.7 | 2 | 12 |
|  | 5 | 933 | 6428.48 | 6.89 | 0.001 | 395.79 | 2527 | 2.7 | 2 | 11 |
|  | 6 | 496 | 5051.01 | 10.18 | 0.001 | 492.95 | 1354 | 2.7 | 2 | 11 |
|  | 7 | 569 | 5169.26 | 9.09 | 0.001 | 317.07 | 1594 | 2.8 | 2 | 15 |
|  | 8 | 583 | 8927.76 | 15.31 | 0.001 | 476.37 | 1574 | 2.7 | 2 | 10 |
|  | 9 | 486 | 6553.37 | 13.48 | 0.001 | 398.72 | 1375 | 2.8 | 2 | 9 |
|  | 10 | 534 | 3905.24 | 7.31 | 0.001 | 434.32 | 1477 | 2.8 | 2 | 10 |
| BFc4 | 1 | 1019 | 14352.62 | 14.09 | 0.001 | 499.04 | 2818 | 2.8 | 2 | 12 |
|  | 2 | 861 | 7904.79 | 9.18 | 0.001 | 415.28 | 2432 | 2.8 | 2 | 11 |
|  | 3 | 796 | 8682.69 | 10.91 | 0.001 | 418.18 | 2153 | 2.7 | 2 | 16 |
|  | 4 | 539 | 6605.65 | 12.26 | 0.001 | 442.01 | 1492 | 2.8 | 2 | 12 |
|  | 5 | 776 | 10870.44 | 14.01 | 0.001 | 479.50 | 2201 | 2.8 | 2 | 15 |
|  | 6 | 476 | 5833.85 | 12.26 | 0.001 | 466.82 | 1278 | 2.7 | 2 | 7 |
|  | 7 | 570 | 4471.35 | 7.84 | 0.001 | 479.70 | 1612 | 2.8 | 2 | 13 |
|  | 8 | 491 | 9272.30 | 18.89 | 0.001 | 495.26 | 1390 | 2.8 | 2 | 12 |
|  | 9 | 541 | 5188.65 | 9.59 | 0.001 | 449.77 | 1478 | 2.7 | 2 | 8 |
|  | 10 | 1236 | 6619.87 | 5.36 | 0.001 | 471.30 | 3371 | 2.7 | 2 | 12 |

The intragenic LD analysis also revealed a higher average |D’| values in the biparental population and the synthetic, compared to the average value observed in the breeding population (0.74 and 0.88 versus 0.67). The biparental population presents an average r^2^ value much lower than the average values observed in the other two populations (0.02 versus 0.13 and 0.14) (Table 4). Regardless of the population, the maximum intragenic |D’| (1,0) was observed for SNPs separated by up to 10.6 kb while most of the higher intragenic r^2^ values (0.7 or greater) were only observed for the closest SNPs (S8 Figure). In regard to the intragenic LD decay, there is evidence of |D’| and r^2^ decay in the breeding population and r^2^ decay in the synthetic (Figure 3). Concerning the intragenic haplotype blocks structure, the general evidence is of a single block of variable size (0.03 to 8.72 kb) with two SNPs (Table 5). Genes Zm00001d018033 and Zm00001d041972 show population differences regarding block size and number of SNPs.

**Table 4.** Intragenic minimum, average, and maximum LD values in each population

| Gene | Population | $\vert D'\vert $ | | | $r^2$ | | |
| --- | --- | --- | --- | --- | --- | --- | --- |
|  |  | Min. | Av. | Max. | Min. | Av. | Max. |
| Zm00001d002654 | Biparental | 0.176 | 0.96 | 1.0 | 0.000 | 0.005 | 0.19 |
|  | Synthetic | 0.003 | 0.60 | 1.0 | 0.000 | 0.159 | 1.00 |
|  | BFc4 | 0.042 | 0.44 | 1.0 | 0.000 | 0.258 | 1.00 |
| Zm00001d004817 | Biparental | 0.028 | 0.81 | 1.0 | 0.000 | 0.004 | 0.06 |
|  | Synthetic | 0.059 | 0.62 | 1.0 | 0.000 | 0.089 | 1.00 |
|  | BFc4 | 1.000 | 1.00 | 1.0 | 0.002 | 0.310 | 0.93 |
| Zm00001d005451 | Biparental | 0.148 | 0.91 | 1.0 | 0.000 | 0.003 | 0.01 |
|  | Synthetic | 0.407 | 0.89 | 1.0 | 0.000 | 0.106 | 1.00 |
|  | BFc4 | 0.057 | 0.51 | 1.0 | 0.000 | 0.211 | 0.97 |
| Zm00001d041972 | Biparental | 0.132 | 0.89 | 1.0 | 0.000 | 0.004 | 0.06 |
|  | Synthetic | 0.263 | 0.79 | 1.0 | 0.000 | 0.191 | 1.00 |
|  | BFc4 | 0.193 | 0.88 | 1.0 | 0.000 | 0.280 | 1.00 |
| Zm00001d052263 | Biparental | 0.236 | 0.85 | 1.0 | 0.000 | 0.011 | 0.06 |
|  | Synthetic | 0.217 | 0.93 | 1.0 | 0.000 | 0.116 | 1.00 |
|  | BFc4 | 0.323 | 0.87 | 1.0 | 0.000 | 0.085 | 1.00 |
| Zm00001d018033 | Biparental | 0.000 | 0.83 | 1.0 | 0.000 | 0.031 | 0.87 |
|  | Synthetic | 0.488 | 0.97 | 1.0 | 0.000 | 0.025 | 0.21 |
|  | BFc4 | 0.137 | 0.77 | 1.0 | 0.001 | 0.070 | 0.46 |
| Zm00001d035760 | Biparental | 0.187 | 0.84 | 1.0 | 0.000 | 0.007 | 0.06 |
|  | Synthetic | 1.000 | 1.00 | 1.0 | 0.007 | 0.007 | 0.01 |
|  | BFc4 | 0.721 | 0.72 | 0.7 | 0.027 | 0.027 | 0.03 |
| Zm00001d036900 | Biparental | 0.000 | 0.76 | 1.0 | 0.000 | 0.093 | 0.88 |
|  | Synthetic | 0.005 | 0.77 | 1.0 | 0.000 | 0.026 | 1.00 |
|  | BFc4 | 0.031 | 0.60 | 1.0 | 0.000 | 0.019 | 0.24 |
| Zm00001d021731 | Biparental | 0.094 | 0.59 | 1.0 | 0.000 | 0.037 | 0.68 |
|  | Synthetic | 0.019 | 0.58 | 1.0 | 0.000 | 0.282 | 1.00 |
|  | BFc4 | 0.193 | 0.57 | 1.0 | 0.001 | 0.248 | 1.00 |
| Zm00001d023810 | Biparental | 1.000 | 1.00 | 1.0 | 0.000 | 0.000 | 0.00 |
|  | Synthetic | 0.026 | 0.76 | 1.0 | 0.000 | 0.093 | 1.00 |
|  | BFc4 | 0.004 | 0.48 | 1.0 | 0.000 | 0.066 | 0.97 |
| Zm00001d025201 | Biparental | 0.097 | 0.84 | 1.0 | 0.000 | 0.004 | 0.06 |
|  | Synthetic | 0.059 | 0.59 | 1.0 | 0.000 | 0.368 | 0.87 |
|  | BFc4 | 0.006 | 0.68 | 1.0 | 0.000 | 0.061 | 1.00 |
| Zm00001d026113 | Biparental | 0.002 | 0.82 | 1.0 | 0.000 | 0.026 | 1.00 |
|  | Synthetic | 0.105 | 0.81 | 1.0 | 0.000 | 0.057 | 0.90 |
|  | BFc4 | 0.015 | 0.52 | 1.0 | 0.000 | 0.073 | 1.00 |

**Figure 3.**
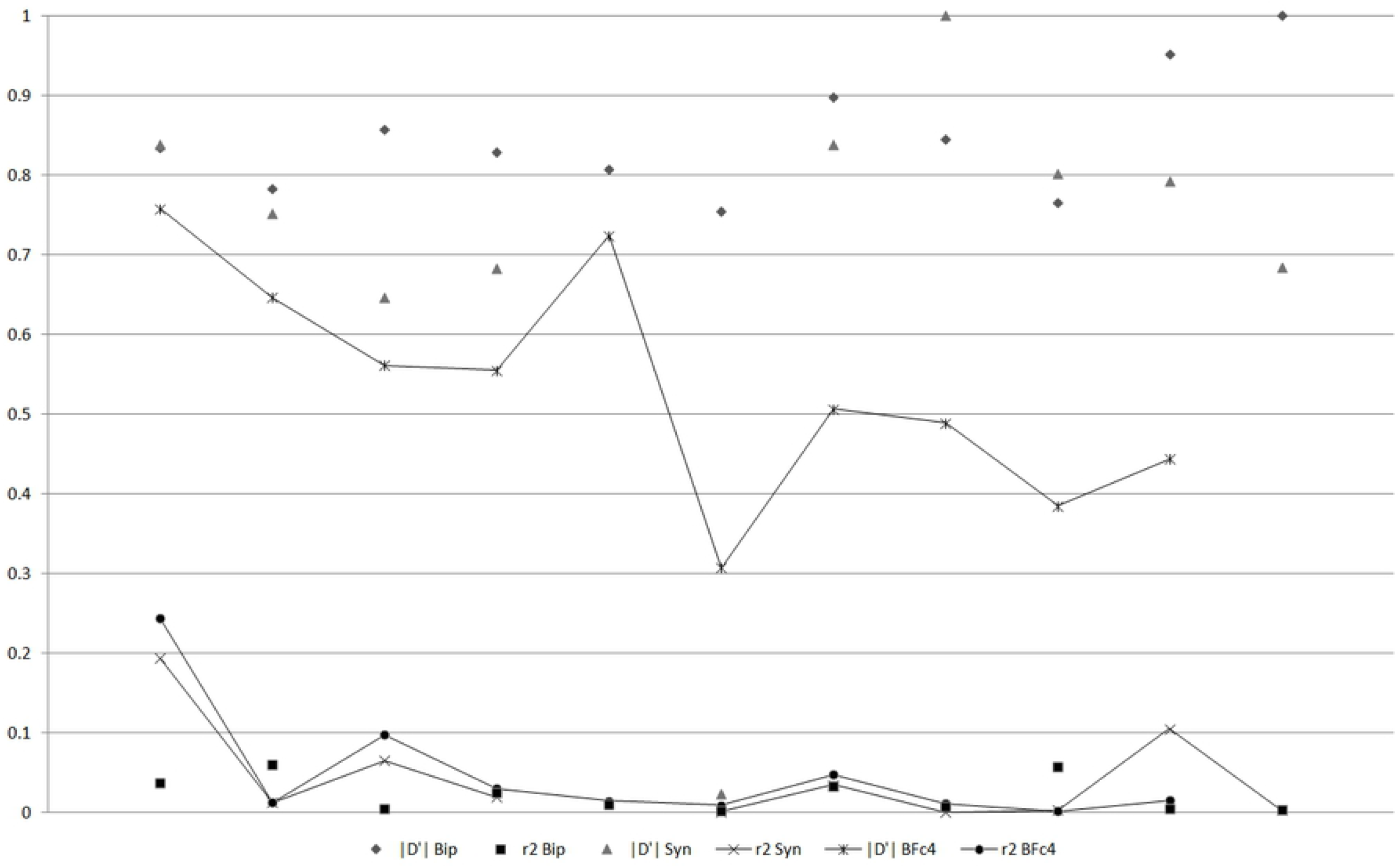
Intragenic LD decay and LD extent concerning SNPs separated by up to 10.6 kb (|D’| and r^2^ average values in intervals of 1 kb).

## Discussion

It is difficult to characterize the LD and haplotype block patterns in two or more unrelated random cross populations based on LD map and two measures of linkage disequilibrium. Based on studies on the LD pattern in human populations, the LD maps demonstrated that the human chromosomes have a pattern of regions of extensive LD (plateaus or cold spots), interspaced with regions of high recombination rate (steps or hot spots) [24, 25]. Both regions are variable in number and length and cold spots show equal (as assumed in this study) or similar LD in LDUs. The hot spots present distinct LDUs. The same pattern was seen in the LD maps of the chromosomes of the biparental population, elaborated under high density as recommended by Pengelly, Tapper (24). To better understand the level of LD in the hot and cold spots, we analyzed two extreme segments of the chromosome 1 LD map, including 30 SNPs. Both segments have similar lengths in LDUs (4.1 and 3.6) and kb (970 and 828). The average |D’| was much greater for the SNPs in the seven cold spots (including three to 12 SNPs), compared with the average value for the SNPs in the 21 hot spots (including two to three SNPs) (0.89 versus 0.29). However, this was not verified via the r^2^ statistic (0.004 versus 0.038).

When comparing populations that share a common origin, have similar effective population size, and did not have face an extreme reduction in size (population bottleneck), because similar allele frequencies the statistics D, D’, and r^2^ should provide a comparable characterization of the LD pattern. If the populations have distinct distributions of the allelic frequencies, D’ can be used for analyzing the recombination history and r^2^ should be the choice if recombination and mutation are important factors affecting the LD [1]. However, in the last two decades most studies on LD in human populations have aimed selecting populations and SNPs (tagging SNPs) for association studies [25, 26]. In general, both |D’| and r^2^ have been used [26, 27] and because their high level of LD, isolated populations have been recommended for association studies [28]. The statistic r^2^ is the most relevant for association mapping because it has a simple inverse relationship with the sample size required to detect association [1]. The use of LD maps and two measures of LD for comparing the popcorn populations provided some contrasting results, but the general evidence is that the synthetic is the population with higher LD. As expected, the lower average |D’| value in the breeding population reflects its recombination history. The synthetic and the biparental population presented greater average |D’| and higher frequency of SNPs with elevated |D’| values because they have no recombination history.

Because the differences regarding molecular marker type and density, sample size, and genome coverage, the comparison of LD values concerning human, domesticated animal, and plant populations should be made with caution even when the studies involve the same species. We were surprised by the low average r^2^ values and the reduced frequency of SNPs with r^2^ values greater than 0.25 (defined as useful LD in some studies) in the popcorn populations. In the study of Yan, Shah (29), involving 632 maize inbred lines and 943 SNPs (density of one SNP each 2,121 kb), the average r^2^ was only 0.009. However, for SNPs separated by up to 100 kb the average was 0.2 (0.03, 0.09, and 0.10 for the biparental, synthetic, and breeding populations, respectively). Even higher LD were reported in the maize NAM (nested association mapping) population [30], and in two biparental and four FPM (four parent maize) populations studied by Anderson, Mahan (9). In general, the average r^2^ values observed in the popcorn populations are also lower than the values observed in cattle and chicken populations (0.1 to 0.8 for SNPs separated by up to 100 kb) [31–33]. The density ranged from 27.8 to 112.3 kb in these three studies. Using a 600K SNP chip (density of one SNP each 6.3 kb), Pardo, Bochdanovits (27) observed a median pair-wise r^2^ averaged across all chromosomes of 0.015 and 0.016 for the Dutch and HapMap-CEU populations, respectively.

The absence of a uniform criterion for defining the LD decay and the LD extent also makes comparing results with human, domesticated animal, and plant populations difficult. Angius, Hyland (25) used LD decay as the distance over which the average LD decreases to half of its maximum value (half-length). They defined LD extent as the distance over which the average LD declines to an asymptotic value. Anderson, Mahan (9) used LD decay as the distance over which the average r^2^ dropped below 0.8, and LD extent as the distance over which the average r^2^ fell below 0.2. Concerning the LD decay, our results showed differences between LD measures and populations. There were slightly differences between chromosomes, but the higher r^2^ decay occurred after 5-10 kb (36 to 73%). Yan, Shah (29) observed a LD decay of 64% after 5-10 kb in an inbred lines panel and the LD reached an approximate asymptotic r^2^ value of 0.01 in the interval of 1-5 Mb (LD extent of 5 Mb). A similar LD extent (5 Mb) was observed in eight breeds of cattle but a comparable LD decay (62%) occurred along 100 kb [34]. From the analysis of segments of one Mb in all chromosomes in Ashkenazi jews, caucasians, and African American populations, Shifman, Kuypers (35) observed LD decays of 17, 21, and 42% along 10 kb, respectively. A similar LD extent of 300 kb occurred in the populations (to reaching an approximate asymptotic r^2^ value of 0.05).

If there is higher LD between QTLs and haplotypes than with individual SNPs, haplotype blocks can provide substantial statistical power in association studies [6] and increased accuracy of genomic prediction of complex traits [36]. Surprisingly, our results evidenced that the number and length of the haplotype blocks and the number of SNPs per haplotype block were proportional to the average r^2^. The criterion of Gabriel, Schaffner (6) appears to provide a reduced number of SNPs per haplotype block. In a study with 235 soybean varieties genotyped by 5,361 SNPs (density of one SNP each 208 kb), Ma, Reif (37) observed six SNPs per haplotype block on average. This is not surprising because the group of varieties correspond to a pure line panel (high LD). In studies with German Holstein cattle and four chicken populations, the average number of SNPs per haplotype block ranged between approximately four to 10 and the mean block length ranged from approximately 146 to 799 kb [31, 32]. Low average numbers of SNPs per haplotype block (approximately 4-5) and reduced average haplotype block lengths (approximately 5-7 kb) were also observed in human populations [6, 27]. However, the size of each block varied dramatically in the study of Gabriel, Schaffner (6), from less than one to 173 kb.

Concerning the low intragenic LD and the minimum size of the haplotype blocks observed in the three populations, we believe that the lower LD for the biparental population is due to crossing two genetically similar high-quality inbred lines. Because there is no information on the LD and haplotype block patterns in the base populations Viçosa and Beija-Flor, we cannot infer that the higher average intragenic r^2^ values observed in the synthetic and breeding population (for 11 of the 12 genes) are due to selection for quality. The characterization of the LD and haplotype block patterns regarding specific chromosomal regions has only been made by human geneticists, generally aiming SNP tagging. From the analysis of SNPs within the HLA region on chromosome 6, Evseeva, Nicodemus (38) observed 18 haplotype blocks in European populations, based on the criterion of Gabriel, Schaffner (6). Furthermore, the LD was slightly lower in southern than northern European populations. Using the same criterion, Nuchnoi, Ohashi (39) observed six and four haplotype blocks across a 472 kb region on chromosome 5q31-33 in Southeast (Thais) and Northeast Asians (Chinese and Japanese) populations. Akesaka, Lee (40) identified two to six blocks in Korean and Japanese populations, depending on the criterion of a LD block, spanning approximately 3 to 47 kb. The median r^2^ value for the five genes in the region ranged from 0.03 to 0.89.

In conclusion, the level of LD expressed by the r^2^ values in the three popcorn populations with different genetic structures - a biparental population, a synthetic, and a breeding population - is surprisingly low, but comparable to some non-isolated human populations. This does not imply that these populations cannot be used for GWAS because there is a fraction of high r^2^ values for SNPs separated by less than 5 kb. The populations are also not excluded for genomic selection because the most important factor affecting this selection process is the relatedness between individuals in the training and validation sets. However, we do not expect a significant advantage from haplotype-based GWAS and genomic selection along generations due to the reduced number of SNPs in the haplotype blocks (2 to 3). The results on LD decay (rapid decay after 5-10 kb) and LD decay extent (along up to 300 kb) are in the range observed with maize inbred line panels. Our most important result is that, similar to the human chromosomes, maize (popcorn is also *Zea mays*, but ssp. *everta*) chromosomes also have a pattern of regions with extensive LD (plateaus or cold spots), interspaced with regions of high recombination rate (steps or hot spots). It should be highlighted, however, that our simple simulated LD map provides evidence that this pattern can reflect regions with differences in allele frequencies and LD level (expressed by D’) and not regions with high and low rates of recombination as evidenced by Jeffreys, Holloway (41), since the simulation process assumes a rate of recombination that is proportional to the distance in cM.

## Acknowledgments

We thank the National Council for Scientific and Technological Development (CNPq), the Brazilian Federal Agency for Support and Evaluation of Graduate Education (Capes; Finance Code 001), and the Foundation for Research Support of Minas Gerais State (Fapemig) for financial support.

## Supporting information

**S1 Table** Gene name, annotation, and chromosome localization, and the number of intragenic SNPs in each population.

**S2 Figure** MAF distribution in the biparental population (a), in the synthetic (b), and in the breeding population (c).

**S3 Figure** LD maps of the populations, by chromosome.

**S4 Table** Minimum and maximum LD values, average distance (kb), and frequency observed in chromosomes by population, concerning SNPs with |D’| and r^2^ values higher than 0.75, in the interval 0.25-0.75, and lower than 0.25.

**S5 Figure** Average |D’| values by chromosome and by distance interval (kb) in the biparental population (a), in the synthetic (b), and in the breeding population (c).

**S6 Figure** Average r^2^ values by chromosome and by distance interval (kb) in the biparental population (a), in the synthetic (b), and in the breeding population (c).

**S7 Figure** Distribution of the haplotype blocks based on the number of SNPs in the biparental population (Bip), in the synthetic (Syn), and in the breeding population (BFc4).

**S8 Figure** Overall intragenic |D’| (a, b, c) and r^2^ (d, e, f) by distance interval (bp) in the biparental population (a and d), in the synthetic (b and e), and in the breeding population (c and f).

## Data availability

The dataset is available at https://doi.org/10.6084/m9.figshare.8277629.v1.

## Author’s contribution

All authors contributed equally.

